# Roles of C/EBPα, GATA2, TGF-β-signaling, and epigenetic regulation in the expression of basophil-specific protease genes

**DOI:** 10.1101/2024.02.24.581851

**Authors:** Ryotaro Tojima, Kazuki Nagata, Naoto Ito, Takahiro Arai, Kenta Ishii, Tomoka Ito, Kazumi Kasakura, Chiharu Nishiyama

## Abstract

In the present study, we analyzed the transcriptional regulation of genes encoding basophil-specific proteases Mcpt8 and Mcpt11 to clarify the molecular mechanisms by which the commitment between basophil and mast cell (MC) is determined. We used bone marrow-derived (BM) cells maintained in the presence of IL-3, in which basophil-like cells and MC-like cells exist. Knock down (KD) and overexpression of a transcription factor C/EBPα showed that *Cebpa* mRNA levels were associated with those of *Mcpt8* and *Prss34* (a gene symbol of Mcpt11). Treatment with trichostatin A (TSA), a histone deacetylase inhibitor, decreased mRNA levels of *Cebpa*, *Mcpt8*, and *Prss34* in BM cells, whereas these mRNA levels were increased by the treatment with an inhibitor of DNA methyltransferase. TSA treatment upregulated mRNA levels of *Gata1* and *Mitf* in BM cells, and *Mitf* KD but not *Gata1* KD increased mRNA levels of *Cebpa* and *Prss34*. Furthermore, *Gata2* KD significantly decreased mRNA levels of *Mcpt8* and *Prss34* without affecting *Cebpa* mRNA level. In addition, mRNA levels of *Gata1* and *Mitf* were increased in BM cells transfected with *Cebpa* siRNA, and decreased in *Cebpa*-overexpressing BM cells. We also found the TGF-β treatment increased mRNA levels of *Mcpt8*, *Prss34*, along with upregulation of *Gata2* and *Cebpa* transcription.

Taken together, we conclude that the transcription of basophil-specific protease genes was positively regulated by C/EBPα, GATA2, and TGF-β signaling with modification of epigenetic regulation.

## Introduction

Basophils and mast cells (MCs), which constitutively express FcεRI in a cell-type specific manner, play important roles in IgE-mediated immune responses, including pathogenesis of allergic inflammation and defense against parasite infection. In addition to their phenotypic and functional similarities, these two lineages can be developed from common progenitors, suggesting that basophils and MCs share a common transcriptional regulatory system in the early stages of the development from hematopoietic stem cells. In contrast, basophils and MCs are clearly distinguished by their distinct profiles of the expression of proteases. That is, basophils specifically express mast cell protease (Mcpt) 8, and 11, whereas mucosal MCs and connective tissue MCs are identified by the expression of Mcpt1 and 2, and Mcpts4-7 and 9, respectively. To elucidate the roles of MCs and basophils in various immune-related diseases, gene-targeted mice have been generated using cell-type specific activation of the *Mcpt* gene promoters. Namely, *Mcpt5*-Cre mice have been used as connective tissue MC-specific Cre mice (1, 2) and the *Mcpt8* gene has been utilized to establish basophil-specific systems (3-5).

We previously demonstrated that expression of the *Mcpt1* and *2* genes in bone marrow-derived MCs (BMMCs) is regulated by the hematopoietic cell-specific transcription factors GATA1 and GATA2 under coordinated regulation with the TGF-β-Smad-axis (6). On the other hand, the regulatory mechanism underlying the expression of basophil-specific protease genes remain unclear. Therefore, in the present study, we analyzed the transcriptional regulation of the *Mcpt8* and *Prss34* (encoding Mcpt11) genes by focusing on the roles of transcription factors, including C/EBPα, MITF, PU.1, Runx1, GATA1, and GATA2, which have been identified as transcription factors regulating the development and gene expression of basophils and/or MCs (7-12). We also evaluated the involvement of epigenetic regulation such as histone acetylation and DNA methylation in the expression of basophil-specific protease genes.

## Materials and Methods

### Mice and Cells

Bone marrow cells isolated from male or female C57BL/6J mice (Japan SLC, Hamamatsu, Japan), which were maintained under specific pathogen-free-conditions, were maintained in BMMC-complete medium, which is RPMI-1640-based and supplemented with 5 μg/mL recombinant mouse IL-3 (#575506 or #575508, BioLegend) as described in previous studies (6, 13).

All experiments using mice were performed following the guidelines of the Institutional Review Board at Tokyo University of Science, and the present study was approved by the Animal Care and Use Committees of Tokyo University of Science: K22005, K21004, K20005, K19006, K18006, K17009, K17012, K16007, and K16010.

### Reagents

DAPI (11034-56, Nacalai Tescue), DMSO (07-4860-5, Sigma Aldrich), 5-aza-2’-deoxycitidine (A2232, Tokyo Chemical Industry), trichostatin A (TSA) (T6933, Fujifilm Wako Pure Chemical), Fc block (clone 93, BioLegend or clone 2.4G2, Tonbo Biosciences), and recombinant mouse TGF-β (5231, Cell Signaling Technology) were purchased from the indicated sources.

The following antibodies were used in flowcytometry, which were obtained from indicated sources: PE anti-mouse FcεRI (clone MAR-1, Tonbo Biosciences, 1:500), FITC anti-mouse CD117 (clone 2B8, BioLegend, 1:500), and APC anti-mouse CD117 (clone 2B8, BD Pharmingen, 1:500).

### Quantification of mRNAs

Total RNA was extracted from cells using the ReliaPrep RNA Cell Miniprep System (#Z6012), RNAzol RT Reagent (#RN190, Cosmo Bio), or ISOGEN (#311-02501, Nippongene). Complementary DNA was synthesized from total RNA by using RevaTraAce qPCR RT Master Mix (#FSQ-201, TOYOBO). Quantitative PCR was performed by a StepOne Real-Time PCR System (Applied Biosystems) with THUNDERBIRD probe qPCR Mix (#QPS-101, TOYOBO) or THUNDERBIRD SYBR qPCR Mix (#QPS-201, TOYOBO) using following primers:

*Cebpa* (Mm00514283_s1, or m*Cebpa*-F; 5’-CGCAAGAGCCGAGATAAAGC-3’ and m*Cebpa*-R; 5’-CGCAGGCGGTCATTGTC-3’); *Gata1* (m*Gata1*-F; 5’-CTAAGGTGGCTGAATCCTCTGC-3’ and m*Gata1*-R; 5’-AGCTACAGAAGATCCTGTCTCC-3’); *Gata2* (Mm00492300_m1, or m*Gata2*-F; 5’-GCCTCTACTACAAGCTGCACAATG-3’ and m*Gata2*-R; 5’-GGGTCTGGATCCCTTCCTTCT-3’); *Mcpt8* (m*Mcpt8*-F; 5’-CAGACCCTGAAGAGGATGTTCCT-3’ and m*Mcpt8*-R; 5’-GGACTCTGTACCCCATATGATTTCC-3’); *Prss34*/*Mcpt11* (mP*rss34*-F; 5’-TGCTTTGTGCTGGGAAGGA-3’ and m*Prss34*-R; 5’-ATGGGCCCCCAGAGTCA-3’); *Mitf* (m*Mitf*-F; 5’-GGAACAGCAACGAGCT-3’ and m*Mitf*-R; 5’-TGATGATCCGATTCAC-3’); *Runx1* (m*Runx1*-F; 5’-CGGTAGAGGCAAGAGCTTCACT-3’ and m*Runx1*-R; 5’-GGCAACTTGTGGCGGATT-3’) and *Spi1* (m*Spi1*-F; 5’-ATGTTACAGGCGTGCAAAATGG-3’ and m*Spi1*-R; 5’-TGATCGCTATGGCTTTCTCCA-3’).

### Plasmid construction

Mouse *Cebpa* full length cDNA was obtained by RT-PCR from total RNA of BM macrophages using forward and reverse primers designed at just upstream of transcription initiation codon and at downstream of termination codon, respectively. After subcloning of PCR products to pCR3.1 (Invitrogen, Thermo Fisher), a plasmid clone carrying complete *Cebpa* cDNA in an appropriate orientation was selected as pCR3.1-mC/EBPα. The DNA fragment isolated from pCR3.1-mC/EBPα by *Nhe*I/*Xho*I-digestion was inserted into *Nhe*I/*Xho*I-digested pIRES2-AcGFP1 (Clontech, Takara) to generate pIRES2-AcGFP1-mC/EBPα.

### Transfection of siRNA and plasmid

We performed transfection of siRNA or expression plasmid by using a Neon Transfection System (Thermo Fisher Scientific). BM-derived culture cells (1.5 x 10^6^) were suspended in 150 μL of T buffer included in the Neon Transfection System 100 μL Kit (Thermo Fisher Scientific). Aliquoted 108 μL T buffer containing suspended cells was mixed with 12 μL of siRNA solution (20 μM for Invitrogen siRNA, and 1 μM for IDT siRNA) or plasmid solution (6 μg plasmid DNA/12 μL TE buffer). After electroporation by a Neon Transfection System set at Program #5 (1700 V, 20 ms, 1 pulse), cells were cultured in BMMC-complete medium without antibiotics.

Following siRNAs purchased from Invitrogen or IDT (TreFECTa series) were used for knockdown experiments: *Cebpa* siRNA (MSS273623), *Gata1* siRNA (MSS236579), *Gata2* siRNA (MSS204585), *Spi1* siRNA (MSS247676), and their GC content-matched appropriate controls from Stealth RNAi siRNA Negative Control Kit (#12935100); *Mitf* siRNA (#108808508), *Runx1* siRNA (#108808494 and #108808497) and the negative control siRNA (#51-01-14-03).

### Flow cytometry

After preincubation in the presence of 1 μg/mL Fc block and 1 μg/mL DAPI for 5 min on ice, cells were stained with antibodies listed in **Reagents** for 15 min on ice. Fluorescence was detected by a MACS Quant Analyzer (Miltenyi Biotec) or a FACSLyric Analyzer (BD Pharmingen). The data were processed with FlowJo software (Tomy Digital Biology).

### Cell sorting

Cells were treated with Fc block and labelled antibodies in the same way as that for flow cytometry and were sorted by using a Cell Sorter SH800 (Sony).

### Determination of methylation degree in genomic DNA

FcεRI^+^/c-kit^-^ (basophil-like) cells and FcεRI^+^/c-kit^+^ (MC-like) cells were isolated from 3 weeks-cultured BM-derived cells by a sorting. A PureLink Genomic DNA Mini Kit (Invitrogen) and a MethylCode Bisulfite Conversion Kit (Invitrogen) were used to purify genomic DNA and to convert methylated cytosine to uracil, respectively. Promoter regions of the *Mcpt8* and *Prss34*/*Mcpt11* genes were amplified by PCR using following primers, which were designed according to the manufacturer’s instructions: *Mcpt8*-methyl-F; 5’-TTTTGGTTGTTTTGGAATTTATTTT-3’ and *Mcpt8*-methyl-R; 5’-AACAAATCCCCTCTCCTATCAATAC-3’; *Prss34/Mcpt11*-methyl-F; 5’-TGGAGGAGTAATGATTGTAGTTAAGG-3’ and *Prss34/Mcpt11*-methyl-R; 5’-ACCAATCAAAACAAATTCACTACCT-3’. The amplified DNA fragments were ligated into T vector pMD19 (#3271, Takara), and whose nucleotide sequences in each plasmid clones were determined.

### Western blotting analysis

Western blotting was performed as previously described (14) using anti-C/EBPα (clone 14AA, #D2205, Santa Cruz Biotechnology) and anti-GAPDH (clone 14C10, #21183, Cell Signaling Technology).

### Statistical analysis

A two-tailed Student’s *t*-test was used to compare two samples. *p*-Values <0.05 were considered significant.

## Results and Discussion

### Expression level of mRNAs for basophil-specific proteases was associated with *Cepba* mRNA levels and was inversely correlated with *Mitf* mRNA levels

We compared mRNA levels of basophils-specific proteases, Mcpt8 and Mcpt11, and transcription factors, C/EBPα and MITF, between 3 weeks (early stage) and 7 weeks (late stage) cultured BM-derived cells, which were maintained in IL-3 supplemented conditions. As shown in **Fig. 1a**, mRNA levels of *Mcpt8*, *Prss34* (encoding Mcpt11), and *Cebpa* were higher in BM-derived cells at 4 weeks, and *Mitf* mRNA levels were significantly higher at 6 weeks. Comparison of mRNA levels between FcεRI^+^/c-kit^-^ (basophil-like) cells and FcεRI^+^/c-kit^+^ (MC-like) cells, which were isolated from 3 weeks-culture BM-derived cells by a sorting (**Fig. 1b**), confirmed that FcεRI^+^/c-kit^-^ cells dominantly expressed mRNAs of *Cebpa*, *Mcpt8*, and *Prss34*, but recessively expressed mRNAs of *Mitf*, *Mcpt1*, and *Cma1* (encoding Mcpt5) compared with FcεRI^+^/c-kit^+^ cells (**Fig. 1c**).

**Figure 1.**
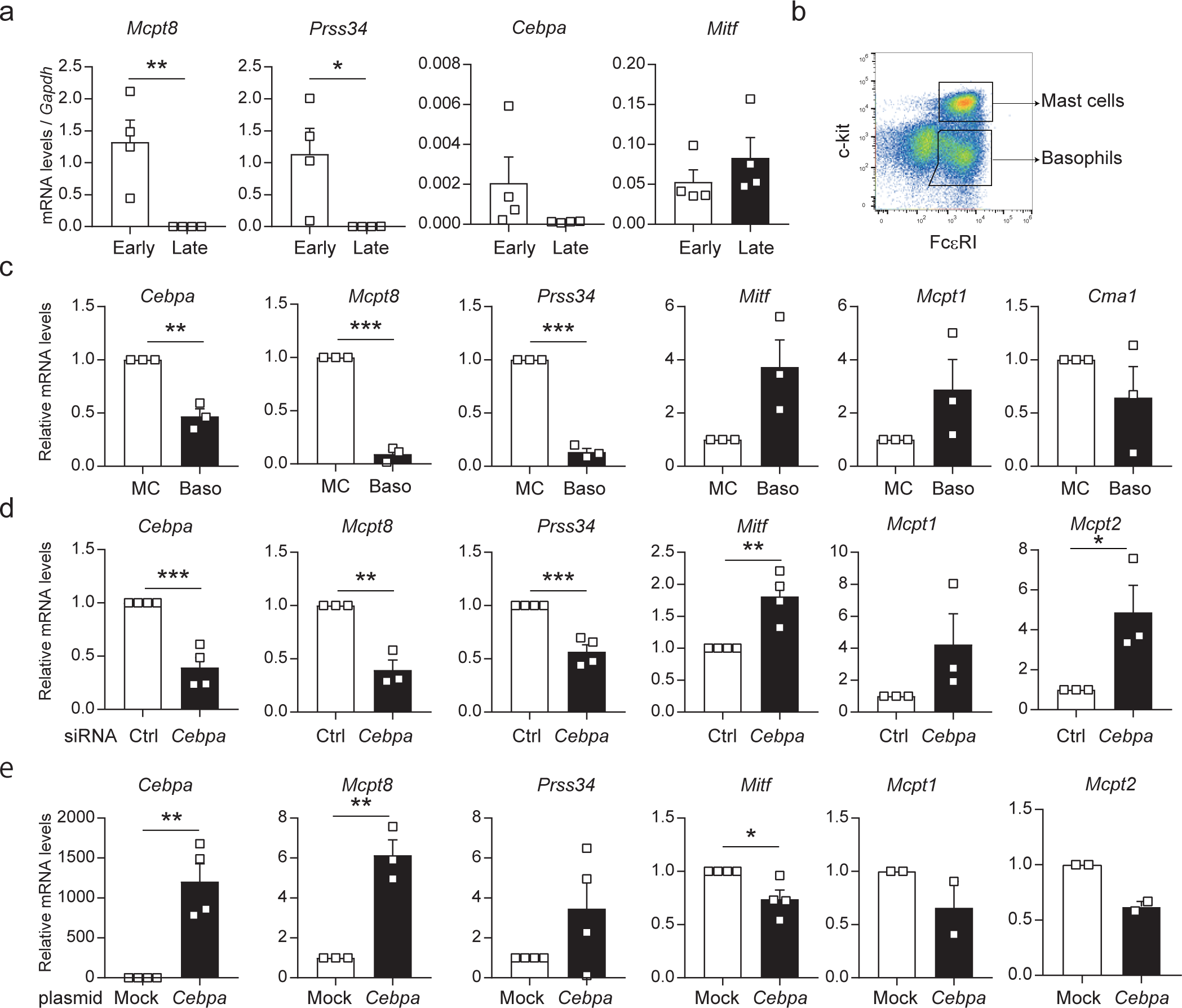
Correlation of expression levels between basophil-specific proteases and related transcription factors. **a.** Messenger RNA levels of *Mcpt8*, *Prss34* (Mcpt11), *Cebpa*, and *Mitf* in 3 weeks- and 7 weeks-cultured BM cells. Early; 3 weeks-cultured BM cells, late; 7 weeks-culture BM cells. **b.** Gating profile of the sorting to isolate MC- and basophil-like cells from 3 weeks-cultured BM cells. **c.** Relative mRNA levels of proteases and transcription factors in MCs and basophils purified from 3 weeks-cultured BM cells by the sorting in Fig. 1b. **d.** Effects of C/EBPα knockdown on mRNA levels in BM cells. BM cells (3 weeks) transfected with *Cebpa* siRNA or its control siRNA were harvested 48 h after electroporation to access RT-qPCR. Ctrl; control siRNA-transfected BM cells, *Cebpa*; *Cebpa* siRNA-transfected BM cells. **e.** Effects of C/EBPα overexpression on mRNA levels in BM cells. BM cells (7 weeks) transfected with expression plasmids were harvested 48 h after electroporation. Mock; BM cells transfected with pCR3.1, *Cebpa*; BM cells transfected with pCR3.1-mC/EBPα. All of the results in **Fig.1a**, **c**, **d**, and **e** are shown as the mean ± SE of data obtained in independent experiments. Significance was determined by tow-tailed unpaired Student’s *t*-test. *p<0.05; **p<0.01; ***p<0.005.

To reveal the roles of C/EBPα in the gene expression of specific proteases and MITF, we performed knockdown and overexpression of C/EBPα, by transfecting siRNA and expression plasmid, respectively. Under the knockdown condition that *Cebpa* mRNA levels were reduced approximately 30% of control, mRNA levels of *Mcpt8* and *Prss34* were significantly decreased, whereas those of *Mitf*, *Mcpt1*, and *Mcpt2* were increased (**Fig. 1d**). In contrast, overexpression of C/EBPα upregulated mRNAs of *Mcpt8* and *Prss34*, and suppressed mRNAs of *Mitf*, *Mcpt1*, and *Mcpt2* (**Fig. 1e**).

These results demonstrate that C/EBPα positively regulates the expression of basophil-specific protease genes, and suppresses the gene expression of MITF and MC-specific proteases.

### Effects of histone acetylation and DNA methylation on the expression of basophil-related genes

To evaluate the effects of epigenetic regulation on basophil-specific genes, we performed experiments using TSA (an inhibitor of HDAC) and 5-aza-2-deoxycytidine (an inhibitor of DNA methyltransferase). When early-stage cells were treated with TSA, the mRNA levels of *Cebpa*, *Mcpt8*, and *Prss34* was significantly suppressed (**Fig. 2a**). The TSA-induced downregulation of *Cebpa* mRNA level reflected to the decrease of C/EBPα protein levels in TSA-treated cells (**Fig. 2b**). On the other hand, 5-aza-2-deoxycytidine treatment tended to increase mRNA levels of *Cebpa*, *Mcpt8*, and *Prss34* in late-phase cells (**Fig. 2c**).

**Figure 2.**
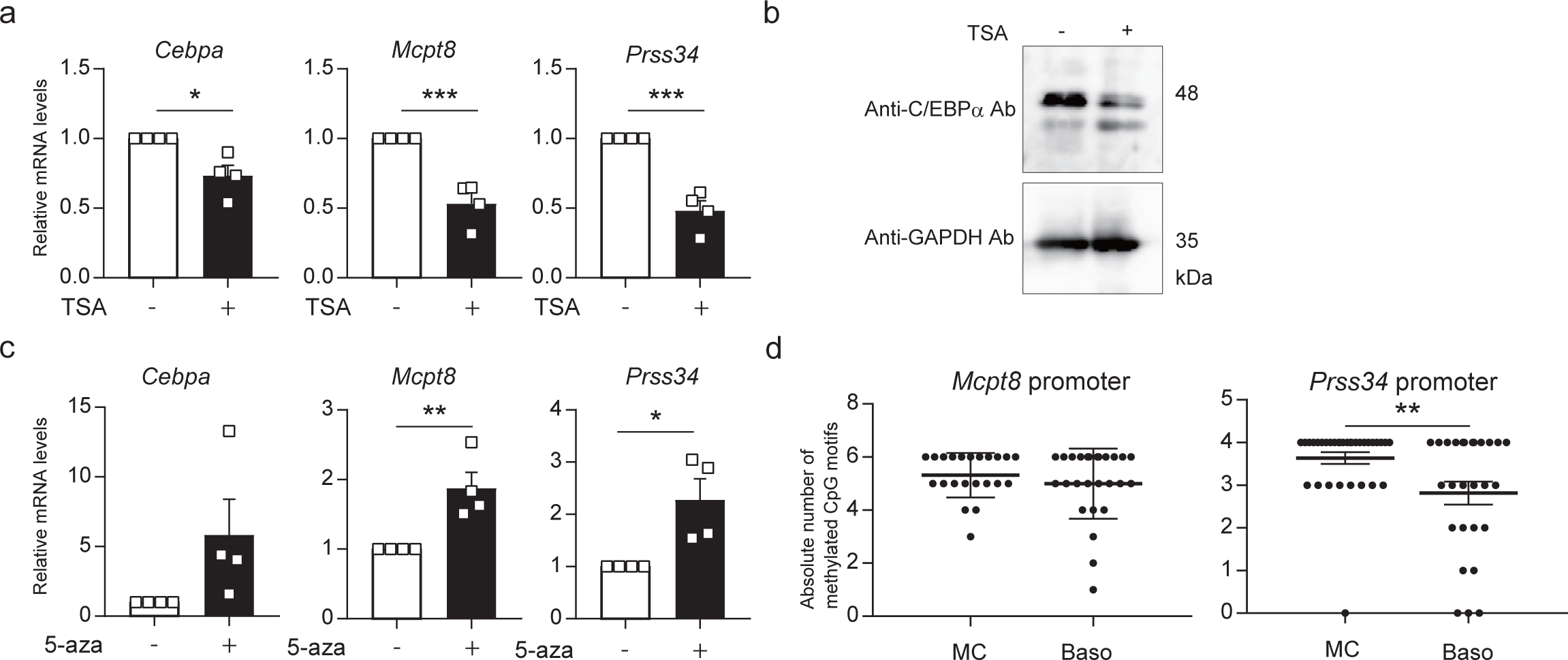
Effects of inhibitors of HDACs and DNA methyltransferases on the expression of C/EBPα and basophil-related genes. **a.** and **b**. Effects of TSA treatment on mRNA expression of *Cebpa*, *Mcpt8*, and *Prss34* (**a**) and C/EBPα protein (**b**). BM cells (7 weeks) were incubated in the presence or absence of 50 μM TSA for 24 h. In a Western blotting, the lysate aliquots of whole cells containing 15 μg of proteins were loaded to each lane (**b**). **c.** Effects of 5-aza-2-deoxycytidine (5-aza) treatment on mRNA levels of *Cebpa*, *Mcpt8*, and *Prss34.* Ten micromole/litter 5-aza-2-deoxycytidine was added 96 h and 48 h before the harvest of 7 weeks-cultured BM cells. **d.** Methylation levels of CpG in the *Mcpt8* and *Prss34* promoters in MCs and basophils. MC- and basophil-like cells were prepared by a same way as in Fig. 1b and c. All of the results in Fig. 2a, and **c** are shown as the mean ± SE of data obtained by independent experiments. Significance was determined by tow-tailed unpaired Student’s *t*-test. *p<0.05; **p<0.01; ***p<0.005.

The knockdown (KD) and overexpression experiments in the present study indicated the correlation between the expression level of C/EBPα and the amounts of transcripts of basophil-specific proteases (**Fig. 1d** and **e**). Above-mentioned results suggest two possible regulatory mechanisms that methyltransferases suppress the transcription of basophil-specific protease genes by methylating the promoters of *Mcpt8* and *Prss34*, or via repressing the expression of C/EBPα by methylating the *Cebpa* promoter. To examine this issue, we quantified DNA methylation degree of these promoters and found that DNA methylation in the *Prss34* promoter was elevated in MC-like cells (**Fig. 2d**).

### Relationship between transcription factors involved in gene expression and development of MC/basophil lineages

Although HDACs generally play roles as negative regulators of transcription, TSA treatment decreased mRNA levels of basophil-related genes (**Fig. 2a**). To reveal the effects of HDAC inhibition on other transcription factors, we determined mRNA levels of MC-related transcription factors in TSA-treated cells. As shown in **Fig. 3a**, TSA treatment significantly enhanced the transcription of *Mitf*, and *Gata1*, but repressed that of *Spi1*. We then investigated the roles of MC-related transcription factors on gene expression of basophil-specific proteases and the gene regulatory relationship between transcription factors by KD experiments. Although *Gata1* KD did not affect mRNA levels of *Cebpa*, *Mcpt8*, and *Prss34* (**Fig. 3b**), *Mitf* KD increased these mRNA levels (**Fig. 3c**). *Gata2* KD significantly reduced mRNA levels of *Mcpt8* and *Prss34* without affecting mRNA level of *Cebpa* (**Fig. 3d**). In addition, it was revealed that the expression of the *Mitf*, *Spi1*, and *Runx1* genes was positively regulated by GATA2 (**Fig. 3d**), and downregulation of PU.1 enhanced transcription of the *Mcpt8* gene (**Fig. 3e**), whereas *Runx1* KD did not affect the expression of basophil-specific protease genes (**Fig. 3f**).

**Figure 3.**
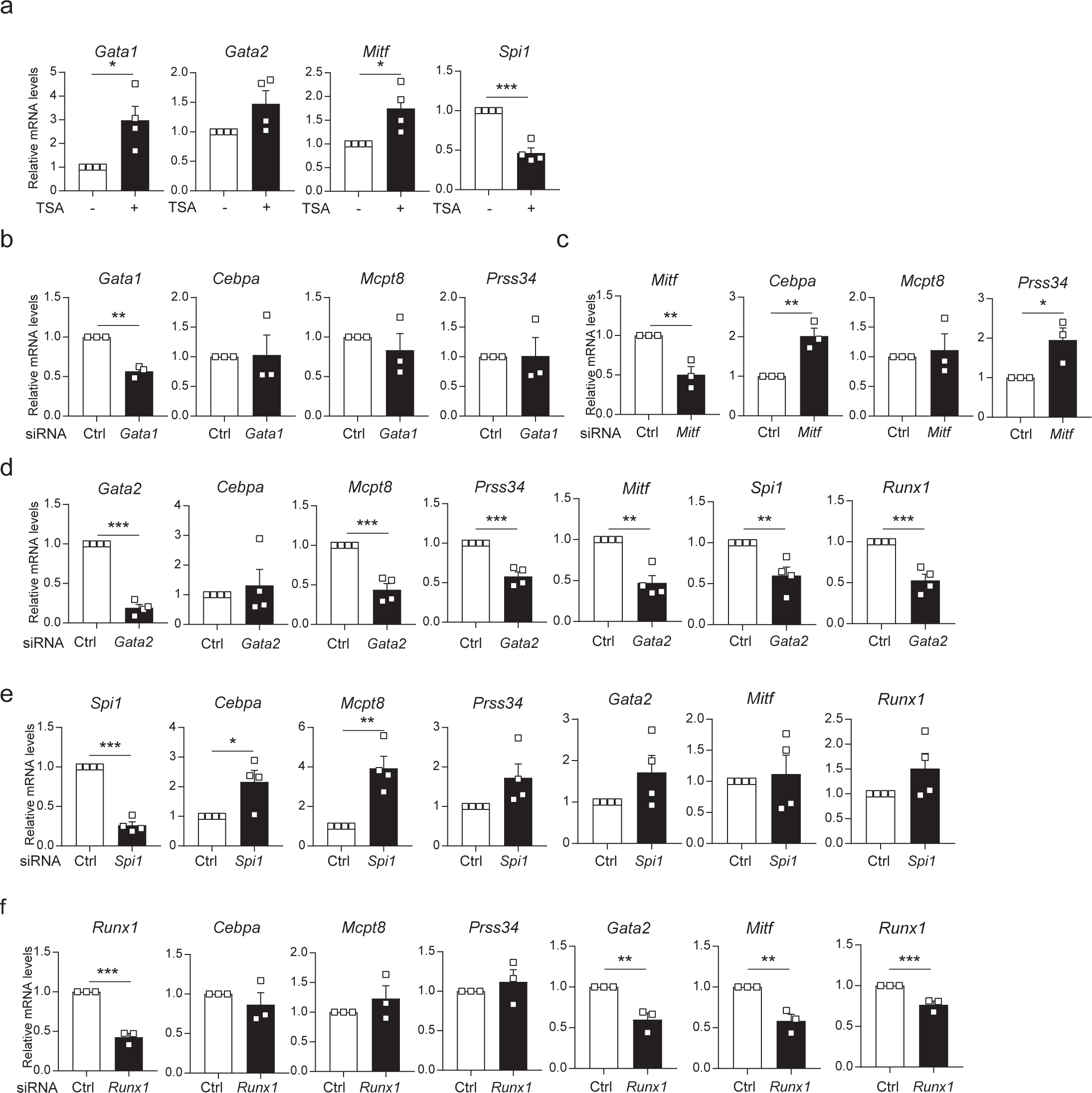
Messenger RNA expression levels of MC/basophil-related molecules in TSA-treated or siRNA-transfected BM cells. **a.** Effects of TSA treatment on mRNA levels of MC-related transcription factors. BM cells (3 weeks) were treated with 50 μM TSA for 24 h. **b**-**f.** Effects of KD of MC-related transcription factors on mRNA levels of *Cebpa*, *Mcpt8*, and *Prss34* in BM cells. KD of *Gata1* (**b**), *Mitf* (**c**), *Gata2* (**d**), *Spi1* (**e**), or *Runx1* (**f**) was performed by siRNA transfection. BM cells (7 weeks) were used for KD of *Gata1*, *Mitf*, *Spi1*, and *Runx1*, and BM cells (3 weeks) were used for *Gata2* KD. All of the results in Fig. 3 are shown as the mean ± SE of data obtained by independent experiments. Significance was determined by tow-tailed unpaired Student’s *t*-test. *p<0.05; **p<0.01; ***p<0.005.

We also analyzed the effects of reduced or enforced expression of C/EBPα on MC-related transcription factors. As shown in **Fig.s 4a** and **4b**, mRNA levels of *Mitf* and *Gata1* were increased and decreased by KD and overexpression, respectively.

**Figure 4.**
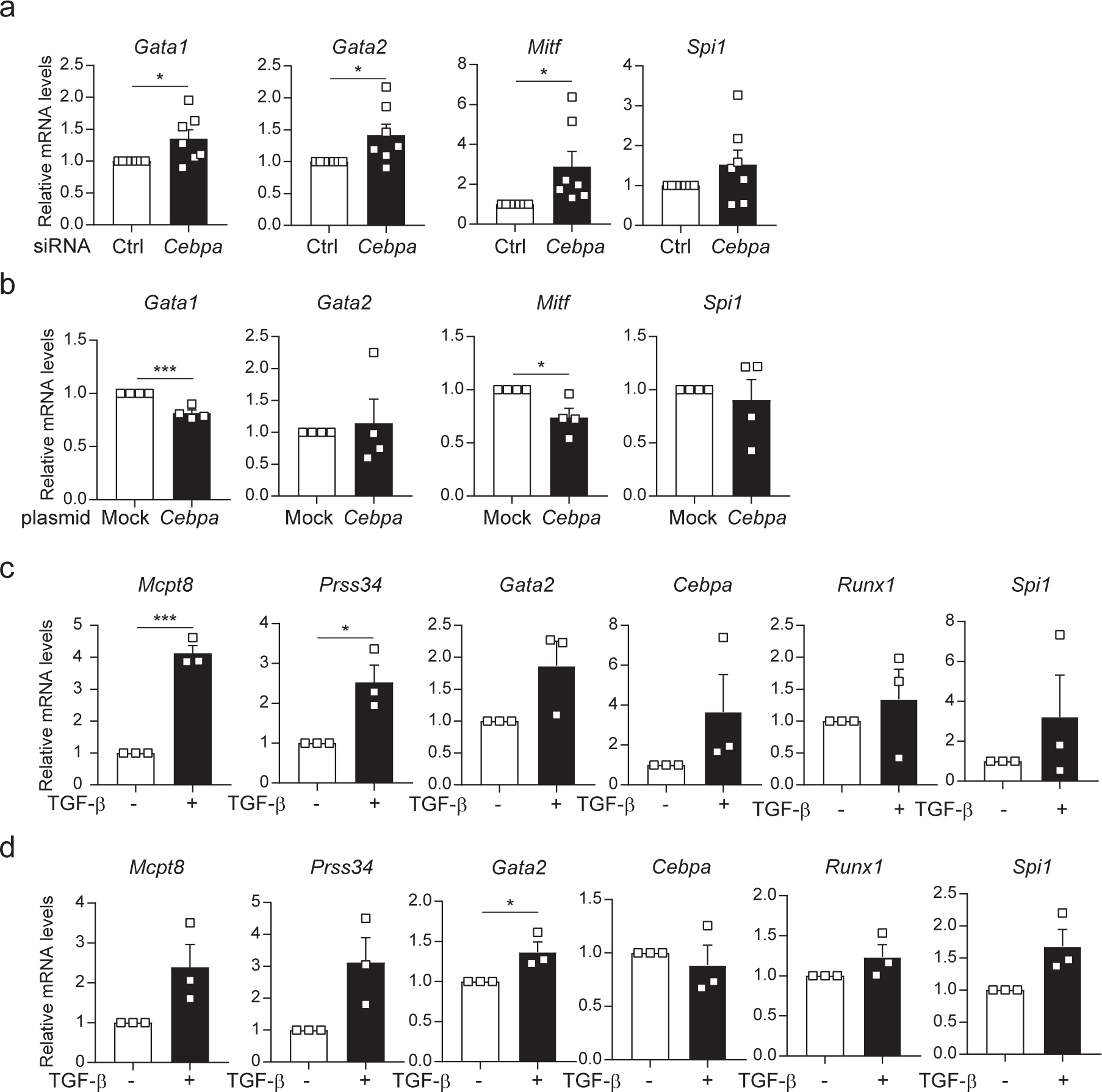
Roles of C/EBPα in the expression of MC/basophil-related transcription factors and effects of TGF-β stimulation on the expression of basophil-specific proteases and related transcription factors. **a.** and **b.** Messenger RNA levels of transcription factors in *Cebpa* siRNA-transfected (**a**) and *Cebpa*-overexpressing (**b**) BM cells. **c.** and **d.** Effects of TGF-β treatment on 7-weeks cultured (**c**) and 3-weeks cultured (**d**) BM cells. All of the results in Fig. 4 are shown as the mean ± SE of data obtained by independent experiments. Significance was determined by tow-tailed unpaired Student’s *t*-test. *p<0.05; ***p<0.005.

From these results, we conclude that GATA2, which is a commonly required for gene expression of basophil/MC lineages, is involved in the expression of basophil-specific protease genes independent from C/EBPα, and that HDAC inhibition accelerated the gene expression of *Mitf* that functions as an antagonistic transcription factor of C/EBPα.

### Roles of TGF-β signaling in the expression of basophils-specific genes

We have previously shown that TGF-β-signaling enhanced transactivation of the *Mcpt1* and *Mcpt2* genes via synergistic regulation by GATA2 and Smads in MCs (6). Although the accelerative roles of TGF-β in the development of mucosal MCs is reported (15, 16), it remains unclear whether TGF-β is involved in the expression of basophil-specific protease genes. Then, we examined the effects of TGF-β stimulation on gene expression of *Mcpt8*, and *Prss34*, and revealed that mRNA levels of *Mcpt8* and *Prss34* in late phase and early phase cells were significantly and tended to be increased by treatment with TGF-β, respectively (**Fig. 4c** and **4d**). Although *Gata2* mRNA level in early BM cells was significantly increased by treatment with TGF-β, mRNA levels of other transcription factors in late (**Fig. 4c**) and early (**Fig. 4d**) phases were not affected by TGF-β treatment. Considering that significant changes were not observed in the levels of *Cebpa* and *Gata2* mRNAs (**Fig. 4c**) in TGF-β-treated BM cells, in which transcripts of *Mcpt8* and *Prss34* were increased (**Fig. 4c**), TGF-β-signaling may enhance the recruitment of C/EBPα and/or GATA2 onto the basophil-specific protease genes.

## Acknowledgments

We thank members of the Laboratory of Molecular Biology and Immunology, Department of Biological Science and Technology, Tokyo University of Science for constructive discussions and technical support.

This work was supported by a Grant-in-Aid for Scientific Research (B) 23H02167 (CN) and 20H02939 (CN); a Research Fellowship for Young Scientists DC2 and a Grant-in-Aid for JSPS Fellows 21J12113 (KN); a Scholarship for Doctoral Student in Immunology (from Japanese Society for Immunology to NI); a Tokyo University of Science Grant for President’s Research Promotion (CN); the Tojuro Iijima Foundation for Food Science and Technology (CN); a Research Grant from the Mishima Kaiun Memorial Foundation (CN); and a Research Grant from the Takeda Science Foundation (CN).

We greatly appreciate the consideration from Dr. Kimihiko Yasuda, Dr. Masako Yasuda, and the late Ms. Yayoi Yasuda.

## Author contributions

R.T., K.N., N.I., and T.A. performed experiments, analyzed data, and prepared figures; K.I., and T.I. performed experiments; K.K. designed research, performed experiments, and analyzed data; C.N. designed research, and wrote the paper.

## References

1. Dudeck, A., Dudeck, J., Scholten, J., Petzold, A., Surianarayanan, S., Köhler, A. et al. (2011) Mast cells are key promoters of contact allergy that mediate the adjuvant effects of haptens Immunity 34, 973–984 10.1016/j.immuni.2011.03.028

2. Scholten, J., Hartmann, K., Gerbaulet, A., Krieg, T., Müller, W., Testa, G. et al. (2008) Mast cell-specific Cre/loxP-mediated recombination in vivo Transgenic Res 17, 307–315 10.1007/s11248-007-9153-4

3. Sullivan, B. M., Liang, H. E., Bando, J. K., Wu, D., Cheng, L. E., McKerrow, J. K. et al. (2011) Genetic analysis of basophil function in vivo Nat Immunol 12, 527–535 10.1038/ni.2036

4. Ohnmacht, C., Schwartz, C., Panzer, M., Schiedewitz, I., Naumann, R., and Voehringer, D. (2010) Basophils orchestrate chronic allergic dermatitis and protective immunity against helminths Immunity 33, 364–374 10.1016/j.immuni.2010.08.011

5. Wada, T., Ishiwata, K., Koseki, H., Ishikura, T., Ugajin, T., Ohnuma, N. et al. (2010) Selective ablation of basophils in mice reveals their nonredundant role in acquired immunity against ticks J Clin Invest 120, 2867-2875 10.1172/JCI42680

6. Kasakura, K., Nagata, K., Miura, R., Iida, M., Nakaya, H., Okada, H. et al. (2020) Cooperative Regulation of the Mucosal Mast Cell-Specific Protease Genes J Immunol 204, 1641-1649 10.4049/jimmunol.1900094

7. Arinobu, Y., Iwasaki, H., Gurish, M. F., Mizuno, S., Shigematsu, H., Ozawa, H., et al. (2005) Developmental checkpoints of the basophil/mast cell lineages in adult murine hematopoiesis Proc Natl Acad Sci U S A 102, 18105-18110 10.1073/pnas.0509148102

8. Tsujimura, T., Morii, E., Nozaki, M., Hashimoto, K., Moriyama, Y., Takebayashi, K. et al. (1996) Involvement of transcription factor encoded by the mi locus in the expression of c-kit receptor tyrosine kinase in cultured mast cells of mice Blood 88, 1225-1233

9. Walsh, J. C., DeKoter, R. P., Lee, H. J., Smith, E. D., Lancki, D. W., Gurish, M. F. et al. (2002) Cooperative and antagonistic interplay between PU.1 and GATA-2 in the specification of myeloid cell fates Immunity 17, 665-676 10.1016/s1074-7613(02)00452-1

10. Mukai, K., BenBarak, M. J., Tachibana, M., Nishida, K., Karasuyama, H., Taniuchi, I. et al. (2012) Critical role of P1-Runx1 in mouse basophil development Blood 120, 76–85 10.1182/blood-2011-12-399113

11. Nishiyama, C., Hasegawa, M., Nishiyama, M., Takahashi, K., Akizawa, Y., Yokota, T. et al. (2002) Regulation of human Fc epsilon RI alpha-chain gene expression by multiple transcription factors J Immunol 168, 4546-4552 10.4049/jimmunol.168.9.4546

12. Maeda, K., Nishiyama, C., Tokura, T., Nakano, H., Kanada, S., Nishiyama, M., et al. (2006) FOG-1 represses GATA-1-dependent FcepsilonRI beta-chain transcription: transcriptional mechanism of mast-cell-specific gene expression in mice Blood 108, 262-269 10.1182/blood-2005-07-2878

13. Oda, Y., Kasakura, K., Fujigaki, I., Kageyama, A., Okumura, K., Ogawa, H., et al. (2018) The effect of PU.1 knockdown on gene expression and function of mast cells Sci Rep 8, 2005 10.1038/s41598-018-19378-y

14. Kitamura, N., Yokoyama, H., Yashiro, T., Nakano, N., Nishiyama, M., Kanada, S., et al. (2012) Role of PU.1 in MHC class II expression through transcriptional regulation of class II transactivator pI in dendritic cells J Allergy Clin Immunol 129, 814-824.e816 10.1016/j.jaci.2011.10.019

15. Miller, H. R., Wright, S. H., Knight, P. A., and Thornton, E. M. (1999) A novel function for transforming growth factor-beta1: upregulation of the expression and the IgE-independent extracellular release of a mucosal mast cell granule-specific beta-chymase, mouse mast cell protease-1 Blood 93, 3473-3486

16. Wright, S. H., Brown, J., Knight, P. A., Thornton, E. M., Kilshaw, P. J., and Miller, H. R. (2002) Transforming growth factor-beta1 mediates coexpression of the integrin subunit alphaE and the chymase mouse mast cell protease-1 during the early differentiation of bone marrow-derived mucosal mast cell homologues Clin Exp Allergy 32, 315–324 10.1046/j.1365-2222.2002.01233.x

